# Multi-source photographic evidence to assess corridor use, crop-raiding behaviour, and body injuries in Asian elephants

**DOI:** 10.1101/2024.11.09.622825

**Authors:** NR Anoop, PK Muneer, M Madhavan, Anikethan Hatwar, T Ganesh

**Affiliations:** Ashoka Trust for Research in Ecology and the Environment (ATREE), Royal Enclave, Srirampura, Bengaluru, Karnataka, 560064, India; School of Social Sciences, Indira Gandhi National Open University (IGNOU), Maidan Garhi, New Delhi, 110068, India; Eco-development committee, Wayanad Wildlife Sanctuary, Tholpetty Range, Kerala, 670646, India

**Keywords:** Asian elephant, multi-source photos, corridor use, landscape connectivity, human-inflicted injuries

## Abstract

This study investigated the use of multi-sourced photographs, such as those from social media, camera traps, and field surveys, for individual identification of male Asian elephants to study their corridor use, crop-raiding behaviour, occurrence of musth, ranging and external body injuries in the Thirunelli-Kudrakote elephant corridor in India’s Western Ghats. A total of 330 images and videos were analysed over 11 years (January 2014 to March 2024), leading to the identification of 27 male elephants from the corridor. Eleven individuals were observed across multiple years, demonstrating their fidelity to the area. Ten elephants were observed engaging in crop-raiding, with four consistently raiding crops within the corridor. The study also assessed the home range of 6 individuals in the corridor and adjacent landscapes based on individual identification. Overall, the study shows that identifying individual elephants across different locations and times provides valuable data and is a practical, cost-effective method for investigating a wide range of questions concerning the ecology and behaviour of Asian elephants. The findings underscore the potential of citizen science initiatives to enhance elephant research across elephant-range countries.

## 1. Introduction

The shrinking and fragmenting habitats of Asian elephants (*Elephas maximus* Linnaeus, 1758), often caused by human encroachment, highlights the special emphasis on understanding how elephants use their habitat. A critical focus within this paradigm is the identification and preservation of ‘elephant corridors’, which have emerged as a paramount management priority to sustain ecological and genetic connectivity while mitigating human-elephant conflicts (Menon et al. 2017; Kanagaraj et al. 2019; Jayadevan et al. 2020). For instance, India is one of the elephant range countries that support around 60% of the global wild population of Asian elephants (Gajah 2010). Elephant habitats in the country are increasingly becoming disconnected, and the species now exists in small, isolated populations, primarily confined to forest areas within fragmented landscapes (Karanth et al. 2010). Given these conditions, the Indian government and international conservation organizations have prioritized identifying and conserving functional elephant corridors to maintain genetic connectivity and support population viability. In India, for instance, 101 identified ‘elephant corridors’ are generally narrow forest pathways connecting larger landscapes, of which 69.3% are regularly used by elephants (Menon et al. 2017). However, many of these critical ‘corridors’ remain vulnerable to various anthropogenic pressures (Mallegowda et al. 2015; Jayadevan et al. 2020).

Individual identification of free-living wildlife species is a commonly used methodological approach for answering a range of research questions in ecology, behaviour, and evolutionary biology, crucial for species management and conservation (Clutton-Brock and Sheldon 2010; Karczmarski et al. 2022). The use of external body features as non-invasive markers through photo-identification stands out as a cost-effective approach for collecting data in individual-based studies within many free-ranging animal populations (Turkalo et al.2013; Wittemyer et al. 2013; Karczmarski et al. 2022). For example, individual adult elephants can be easily distinguished by their unique morphological features, allowing for effective photographic sampling (Goswami et al. 2007; Bedetti et al. 2020). Hence, individual recognition through image-based identification has been increasingly used to estimate various population parameters and to study the behaviour of elephants at individual and population levels through space and time, such as density, distribution, and behaviour (Mumby and Plotnik 2018; Srinivasaiah et al. 2019; Chan et al. 2022). Such methods are particularly feasible in the Indian context for several reasons, including (1) strict legal protection of elephants makes obtaining permissions for radio-collaring challenging and costly; (2) high human population density, including in rural and forest-fringe areas, increases the likelihood of elephant sightings and facilitates photographic data collection; and (3) local communities possess valuable traditional knowledge about elephant movements and behavior, which can be leveraged to support monitoring efforts.

Considering the general knowledge gap on ‘corridor use’ by Asian elephants, the aim of this study was to test the efficiency and reliability of multi-source photographs of male Asian elephants to understand the use of the Thirunelli-Kudrakote ‘elephant corridor’ in the Wayanad elephant Reserve of the Western Ghats (See Menon et al. 2017), their site-fidelity to the area and crop-raiding behaviour. This study focused on male elephants because they are easily identifiable in photographs, particularly due to their larger size and distinct physical characteristics. Males are more susceptible to poaching and are disproportionately involved in crop raiding compared to females in the study area (See Anoop et al. 2023a); hence, understanding the behaviour of the male component of the population is crucial for gaining insights into the ultimate causes of human-elephant interactions and potential for applied conservation research. In this study, we show how opportunistically collected multi-sourced photographs of elephants can be used to study their use of the ‘corridors’, long-term fidelity to the area, and movement to and from adjacent landscapes.

## 2. Materials and methods

### 2.1 Study site

The study was conducted in the Thirunelli-Kudrakote’ elephant corridor’ (located between 11° 53’ 9”-11° 54’ 44” N and 76° 0’ 19”-76° 3’ 55” E) and the Wayanad Sanctuary and Begur Range of North Wayanad division in the Wayanad district of Kerala state (See Fig. 1). This ‘corridor’ is a vital forest connectivity in the Brahmagiri-Nilgiri-Eastern Ghats elephant landscape that holds the largest breeding population of Asian elephants globally (Menon et al. 2017). Extending over a length of 6 km and a width ranging from 1 to 1.5 km, this ‘corridor’ serves as a crucial forest connectivity between the wet evergreen forests and high-altitude grasslands of the Brahmagiri hills in the west and the moist and dry deciduous forests of the Nilgiri Biosphere Reserve in the east. The predominant vegetation types include moist deciduous forest, teak and eucalyptus plantations. The forest has been subjected to intense anthropogenic disturbances such as forest fires, cattle grazing, and NonTimber Forest Products (NTFP) collection (Anoop and Ganesh 2020; Anoop et al. 2023b).

**Fig. 1.**
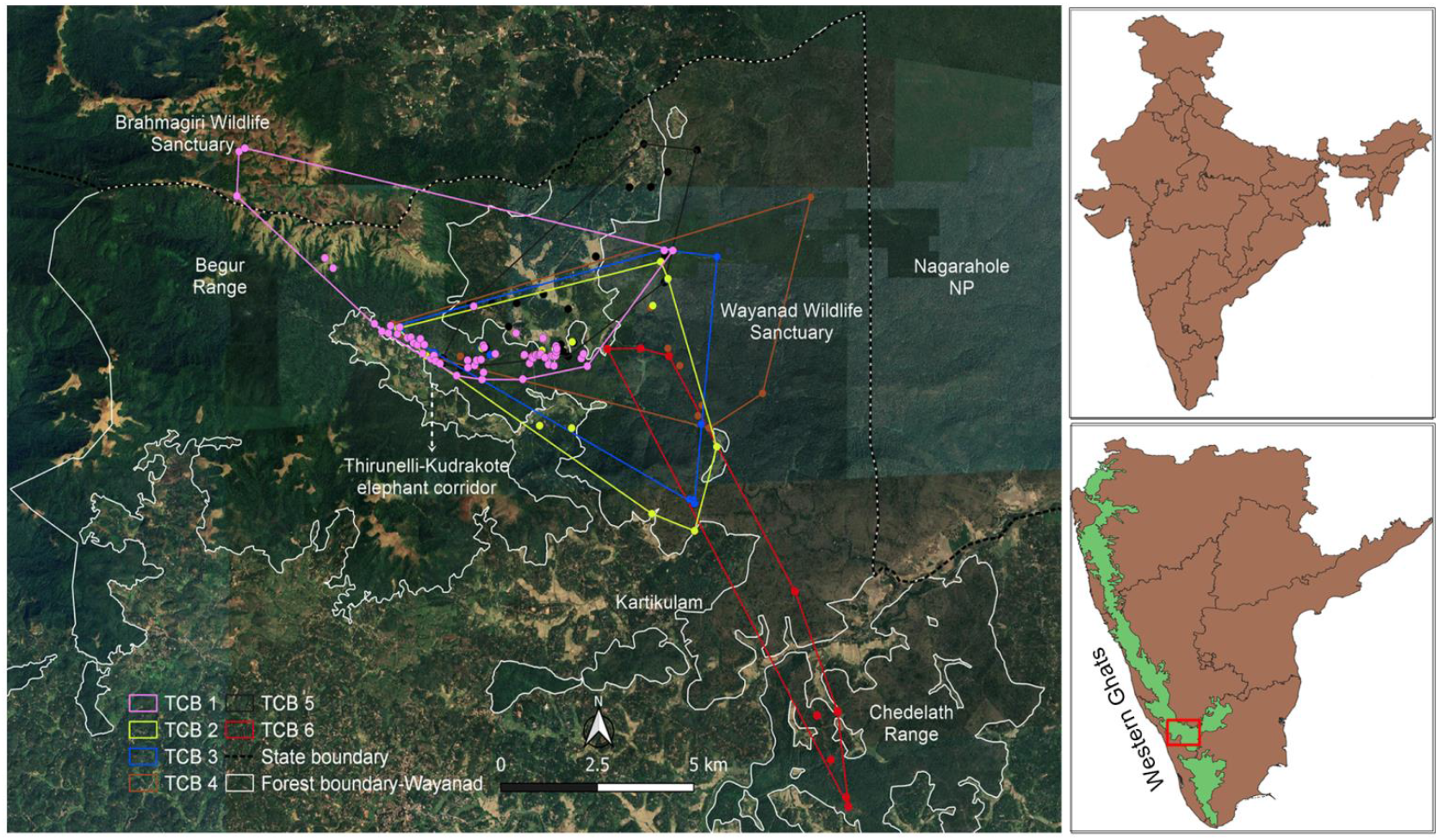
Map showing the Thirunelli-Kudrakote elephant corridor and its connected forest regions, including the Brahmagiri hills to the west and the Tholpetty range of the Wayanad sanctuary to the east. The map also illustrates the minimum convex polygon of six individuals based on available sightings (TCB = Thirunelli Corridor Bull; refer to Figure 2).

The corridor is intersected by two public roads. The land use around the corridor is a heterogeneous matrix of human land use, including plantations with various cash crops and open agriculture such as paddy and plantain in the wetlands. Coffee is a major crop in the area, usually intercropped with pepper, areca nut, coconut, and plantain. The major threats to the corridor include traffic along the roads, the spread of invasive plants such as *Senna spectabilis*, fire and livestock grazing (Anoop et al. 2023a). Human-elephant conflict is severe in the corridor, where male elephants are mostly responsible for crop-raiding (Anoop et al. 2023a).

### 2.2 Method

### Web image search

From January 2014 to March 2024, elephant photographs were collected through manual searches on platforms like Facebook, Instagram, and Google image search using terms like “elephant,” “Thirunelli,” “Thirunelli-Tholpetty,” and “Thirunelli-Kudrakote elephant corridor”. Additional images were procured from local hotels and resorts’ websites, YouTube channels, photographers, forest department officials, and residents. The photograph locations were verified using associated web pages, landmarks, or communication with photographers. Duplicate photos were removed through a final verification process that checked for repetitions by the same photographer or on the same date (See Kannan et al. 2023). **Photographing elephants and camera trapping:** Between 2018 and 2022, two to three observers walked or drove a motorbike along the trails and roads and photographed elephants opportunistically from the study area. Surveys were conducted in a non-systematic manner, with the objective of achieving broad coverage and recording as many different individuals as possible. Ideally, photos were taken from both the right and left flanks of the individuals, although this was not always possible as elephants were concealed within dense understory or moved away before both photographs could be captured. Each encounter was carefully documented, noting the date, time, GPS location, age, and distinctive features of the elephants. At night, we visited villages that frequently experienced crop raiding to photograph and identify the individuals involved in crop-raiding. To effectively target our efforts, we regularly gathered information about crop-raiding incidents from both local villagers and forest department officials. This collaboration allowed us to stay informed about which villages were experiencing conflicts at any given time and to identify patterns of elephant movement related to these incidents. Three automatic camera traps (one Cuddeback and two Hawk ray Trail Cameras) were deployed along the paths frequently used by elephants to enter villages: November-December 2022, June-July 2023, and September 2023 to March 2024. Cameras traps mounted at optimal heights, captured photos and 20-second videos continuously. Traps were checked fortnightly to retrieve data and replace batteries.

### 2.3 Image analysis

Photos and videos were rated as poor, good, or excellent based on focus, framing angle, and visible body proportion. Only images allowing individual elephant identification were analysed. Identification was based on distinct external features such as ear tear patterns, nicks and holes, tusk shape and size, tail morphology, body scarring, deformities, and injuries (Goswami et al. 2007; Vidya et al. 2014). Each individual was assigned a unique code (e.g., ‘TCB 1’ for Thirunelli Corridor Bull 1), with photos and details stored in a computer (see Fig. 2). Capture history was recorded for each individual, with matches verified by at least one additional observer to minimise false positives. The range of six individuals with ample captures and movement across different landscapes was delineated using minimum convex polygons (locations pooled across years and seasons) with convex hull geometry in QGIS version 3.4 (Fig. 1). MCP estimates an animal’s home range by connecting 100 % of the location points that make up the boundary of the range and calculating the area of the resulting polygon. Even though this method is strongly influenced by outliers, this technique is the simplest and most comparable method for calculating home ranges and has been used by several studies to estimate the home range of elephants (See Fernando et al. 2008; Alfred et al. 2012).

**Fig. 2.**
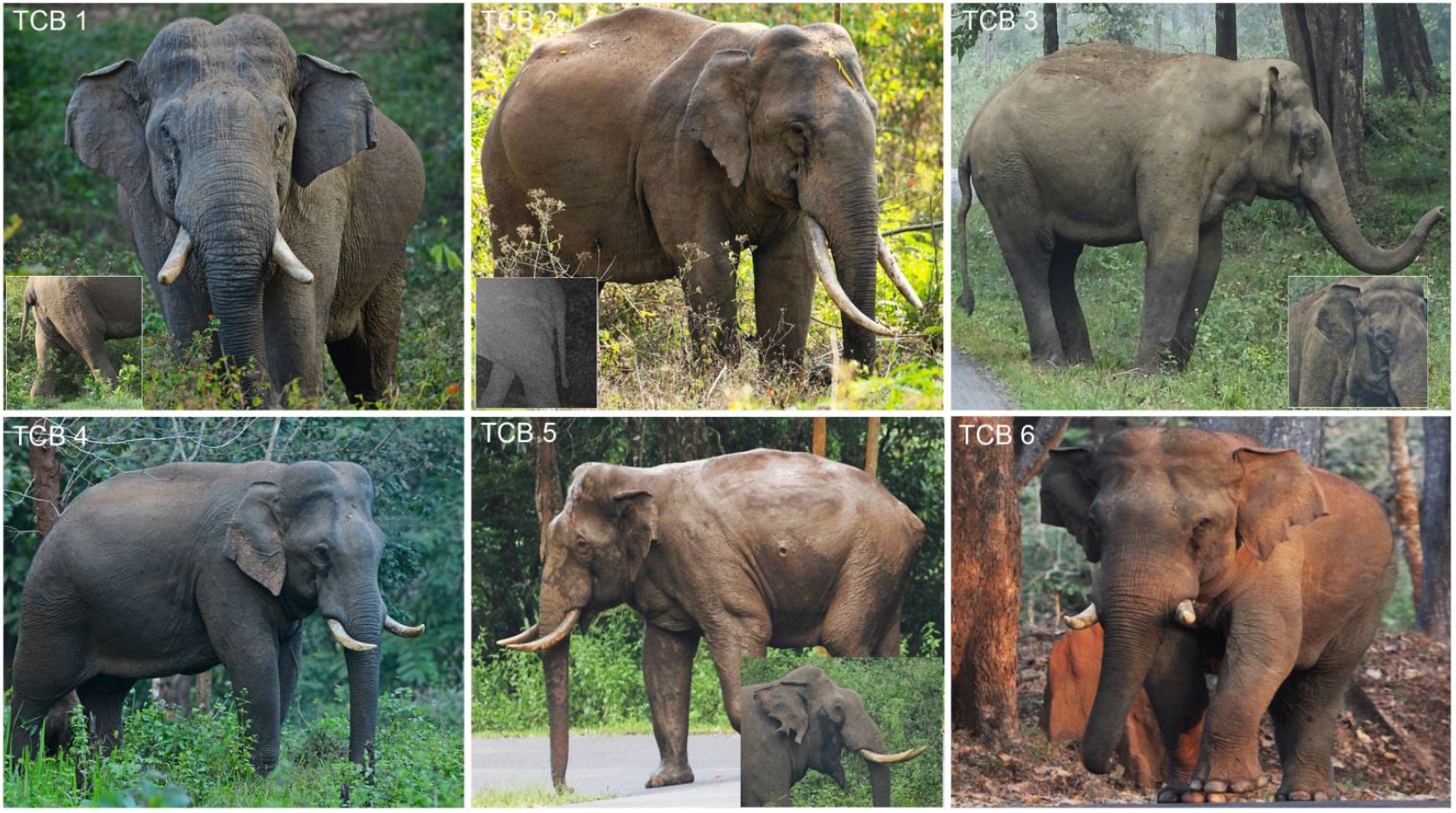
Commonly used morphological features for individual identification of elephants during the study (see Goswami et al., 2007; Vidya et al., 2014): TCB 1 - 1) Ear: primary fold forward and secondary fold backward, 2) small hole on the left ear, 3) lump on the trunk, 4) divergent tusks, 5) broken tail (middle); TCB 2 - 1) Ear: primary fold straight, secondary fold backward, small holes on the left and right ears, 2) convergent tusks, left higher, 3) long tail with brush; TCB 3 - 1) Ear: primary fold slightly forward, secondary fold backward, 2) crooked/bent tail, 3) small hole on the right ear; TCB 4 - 1) Ear: primary fold slightly forward, secondary fold backward, small tear on the right ear, 2) convergent tusks, left higher, 3) long tail with brush; TCB 5 - 1) Ear: primary fold forward, secondary fold backward, part missing from the right ear, 2) broken tail at the base, 3) divergent upward tusks, lump above the right foreleg and below the shoulder blade; TCB 6 - 1) Ear: primary fold forward straight, secondary fold backward, 2) semicircular part missing from the left ear, 3) divergent upward tusks.

## 3. Results and discussion

A total of 330 images and videos were analysed over ten years, leading to the identification of 27 male elephants from the corridor. Among these, 11 individuals were observed across multiple years, demonstrating their fidelity to the area. In contrast, 4 individuals were documented only once (see Appendix A: Fig. S1 & Table S1). The study also assessed the home ranges of 6 individuals in the corridor and the adjacent landscapes based on individual identification (Fig. 1). 10 elephants were observed engaging in crop-raiding, with 4 of them (TCB1, TCB2, TCB5, and TCB12) consistently raiding crops within the corridor and adjacent Tholpetty Range (Appendix A: Table S1). There was a notable congregation of male elephants during the jackfruit (May-September) and paddy seasons (June-December) in Wayanad. During the day, most of the crop-raiding elephants stay close to human habitations and move to farmlands at night to raid crops (Anoop et al. 2023). Consequently, while elephant corridors in India are crucial for landscape connectivity, they can also become conflict hotspots. This is attributed to the irregular forest boundaries with extensive anthropogenic edges; forest areas can be elephant refuges that offer resources, hence allowing elephants to remain close to human habitations during the day without threats and availability of food crops in nearby farmlands. Furthermore, the degradation of corridors exacerbates human-elephant conflicts.

### 3.1 Corridor ‘use’ and behaviour of elephants

Elephants using the corridor demonstrate individual variations in behaviour, age, movement, musth season personality, interaction with people in the area, and crop-raiding tendencies (See Appendix A: Table S1). To provide diverse contexts regarding corridor ‘use’ by different individuals, we focus on six individuals with a maximum number of sightings (See Appendix A: Table 1) who moved to different landscapes during musth and out of musth season (See Fig. 2). TCB 1: An adult bull known locally as “Kannan” is bold yet not aggressive and is frequently seen along the road that runs parallel to the corridor. During non-musth, he frequents the corridor between Thirunelli and Karamat village. During musth, he moves to the high-altitude grassland-evergreen forests of the Brahmagiri hills (See Fig. 1). TCB 1 often raids crops, with heightened activity between May and August, before musth begins. TCB 2 is an adult bull and one of the persistent crop raiders in the area, involved in raiding throughout the year. He is bold and ‘aggressive’ and is found in a large area from the south-near Kartikulam town to the western parts of the corridor (Thirunelli). TCB 3, an adult makhna, uses areas from the western parts of Thirunelli in the corridor and north, central, and southern parts of Tholpetty Range during musth and non-musth seasons. This bold individual exhibits aggression towards people and vehicles during musth and engages in crop-raiding, though less frequently than TCB1, TCB2, and TCB5. There are no recorded sightings of TCB 3 in the Brahmagiri hills. TCB 4 is a calm adult bull mostly found in the northern parts of the Tholpetty range but occasionally appears in the corridor. He travels to the Tholpetty and corridor areas during musth and is rarely seen during non-musth periods. TCB 5, known locally as “Zerotail” or “Murivalan,” is an adult bull that travels the corridor and northern parts of the Tholpetty Range during musth and non-musth periods. He is commonly seen along the main road and has not been recorded in the Brahmagiri hills. Zerotail, known for his aggression towards humans, is a persistent crop raider in the region. He raids crops in the Tholpetty, Appapara, Karamat, Panavally, and Aranappara areas. Additionally, Zerotail stands out as one of the most injured individuals. TCB 6 is a bold adult bull primarily inhabiting the Chedelath Range on the right bank of the Kabini River. During musth, he migrates to the Tholpetty Range and corridor areas. He is known for raiding crops in the Chedelath Range but has not been observed raiding crops in the corridor area.

Due to the expansion of human settlements, the forest connectivity between the Chedelath and Tholpetty ranges has been lost. As a result, the elephants travel through farmlands to reach the Tholpetty range and the corridor. These observations underscore the value of photo-capturing elephants randomly to understand elephant movements across both intact and transformed landscapes. The study was limited to Kerala state, restricting the monitoring of elephant movements across neighbouring states. Consequently, the actual home range of these elephants is likely to be larger than the currently estimated area (See Fig. 1).

Regarding the movement and range of elephants, the current method has disadvantages compared to the technologically advanced radio-telemetry methods that can collect fine-scale spatiotemporal location data from a distance without visual contact with the animals being studied. Although radio telemetry offers fine-scale spatiotemporal data, it is logistically and financially challenging, making large-scale monitoring difficult, especially in developing and underdeveloped countries like India (Miller et al. 2010; Habib et al. 2014). Despite these constraints, visual identification of elephants at different locations and times can yield valuable insights into movement, site fidelity, use of functional corridors, and crop-raiding behaviour (See Desai 1991; Fernando et al. 2011). Therefore, we recommend implementing individual-level identification and monitoring of elephants across all elephant habitats to aid in the conservation of the species.

### 3.2 Understanding behaviour individuality and conflict mitigation

Human–elephant conflict has recently become a topic of major concern in elephant conservation (See Shaffer et al. 2019). Recently, there has been a growing emphasis on incorporating the behavioural individuality of elephants into conflict mitigation (See Mumby and Plotnik 2018). Our previous study found that only a small proportion of male elephants are responsible for most of the crop-raiding incidents in Wayanad (See Anoop et al. 2023a). We found variability in behaviour among the individuals using the corridor, such as their crop-raiding behaviour, aggression towards people, boldness, site-fidelity, musth-season, and use of specific paths to enter villages (See Appendix S1: Table S1). This highlights a detailed understanding of individual variation in life history, personality and behaviour of crop-raiding individuals for the applied conservation. For instance, one of the approaches to reduce conflict in elephant range countries is capturing crop raiders and keeping them in captivity or translocating to other landscapes (Saaban et al. 2011; Fernando et al. 2012). However, this study shows that crop-raiding elephants use different areas of the landscapes during their musth and non-musth seasons. For instance, TCB 1 moved to Brahmagiri during musth seasons from the corridor area, and most of the other individuals were present in the corridor and areas to the east of the corridor during musth and non-musth seasons. This highlights the need to understand the personality differences, home ranges, associations with other individuals, and patterns of gene flow within fragmented habitats before removing elephants from any landscape for conflict mitigation. Identifying crop raiders and non-crop raiders is important for mitigating conflict through radio telemetry or developing individual-specific conflict mitigation methods. Also, crop-raiding is a learned behaviour; hence, managing persistent crop raiders will help prevent the recruitment of new crop raiders in elephant landscapes.

Photographing elephants in their natural habitat is a valuable method for studying various behavioural aspects, such as musth in male elephants characterised by increased sexual and aggressive behaviours. This condition can be influenced by factors such as body injuries, food availability, and age, affecting its timing, duration, and other characteristics in different individuals (Poole and Moss 1981; Keerthipriya and Vidya 2019). A total of seven individuals were observed in musth, with all occurrences of musth taking place in the post-monsoon (September and December) season (Appendix S1: Table S1). Randomly taken photos or camera trap images can also help identify ‘wanderers’ and ‘homers’ in regions where elephant translocations are a conservation strategy (Fernando et al. 2012). For instance, TCB 18 (see Appendix S1: Fig. S1) was tranquilised and relocated from Hassan district to Bandipur Tiger Reserve, a distance of about 130 km. This individual was later photographed in the corridor and subsequently moved to Mananthavady town, where it unfortunately died due to improper management. Early information on the movement of translocated ‘problem-elephants’-such as TCB18’s to Wayanad from other landscapes can be crucial for their immediate management, including capturing or herding them back into the forest before they enter human habitation. Photographs of elephants are also helpful for understanding their dispersal processes into new landscapes.

### 3.3 Conflict-induced scars and injuries in free-ranging elephants

Studies have demonstrated that free-ranging elephants in Africa (Obanda et al. 2008; Mijele et al. 2013) and Asia (LaDue et al. 2010) frequently sustain injuries, with adult males involved in crop-raiding being particularly susceptible. Between 2020 and 2023, eight elephants in our study area sustained human-inflicted injuries, some of which were life-threatening, such as wounds from sharp objects like arrows. Persistent crop-raiding elephants (TCB1, TCB2, TCB5, and TCB18; see Appendix S1: Table S1) experienced a higher incidence of injuries. As expected, these injuries were most common during the paddy and jackfruit seasons, with the legs and body being the most commonly affected areas. However, this study did not observe leg injuries from snares, which are common in Africa (see Obanda et al. 2008). Although injuries among free-ranging elephants are prevalent due to natural and human-induced factors, their impact on elephant health in India has not been thoroughly investigated. Moreover, there is a lack of established mechanisms for identifying and treating injured elephants in the wild. We recommend monitoring and treating both human-induced injuries (e.g., from conflict, snares, and vehicle collisions) and natural injuries (e.g., from fights) in the field, as some can be life-threatening (Mijele et al. 2013).

## 4. Conclusion

Overall, this study demonstrates that photographs of elephants from various sources and field observations are invaluable for understanding landscape connectivity, distribution, behaviour, and injuries among free-ranging Asian elephants. In Africa and Asia, elephants garner significant interest from tourists, researchers, and photographers, making them one of the primary attractions on these continents. Also, the widespread use of smartphones has created new opportunities in the field of citizen science. Developing mobile-based applications where anyone can contribute images of elephants with additional details can help monitor elephant populations, primarily their movement on large spatial scales. This approach can be utilised to promote long-term data collection on elephants through well-planned and organised citizen science programs. We also recommend that the forest departments create photo libraries of elephants in their parks and regularly monitor their movements, corridor usage, crop-raiding behaviour, and physical injuries. This data is essential for understanding individual variations in behaviour and cognition, which are increasingly acknowledged for applied conservation.

## Acknowledgements

We extend our sincere gratitude to the Chief Wildlife Warden and the Principal Chief Conservator of Forests, Kerala Forest Department, for granting the necessary permits for this research. Our heartfelt thanks go to the officers and field staff of the Forest Department for their invaluable assistance with logistics. We are particularly grateful to Abdul Gafoor, Deputy Range Officer, and foresters Sunil and Unni for their support during the fieldwork and for providing elephant images and videos. We also thank Nidheesh and Anurag Varakil for contributing elephant images. We acknowledge the financial support from The Rufford Foundation, United Kingdom, and the equipment support by Idea Wild. Finally, we thank Arjun Kannan for reviewing this draft and offering valuable comments to improve the quality of the manuscript.

