## supplementaryfile1 for "Multi-source photographic evidence to assess corridor use, crop-raiding behaviour, and body injuries in Asian elephants"

**Figure S1.** Male elephants identified from Thirunelli-Kudrakote' elephant corridor'

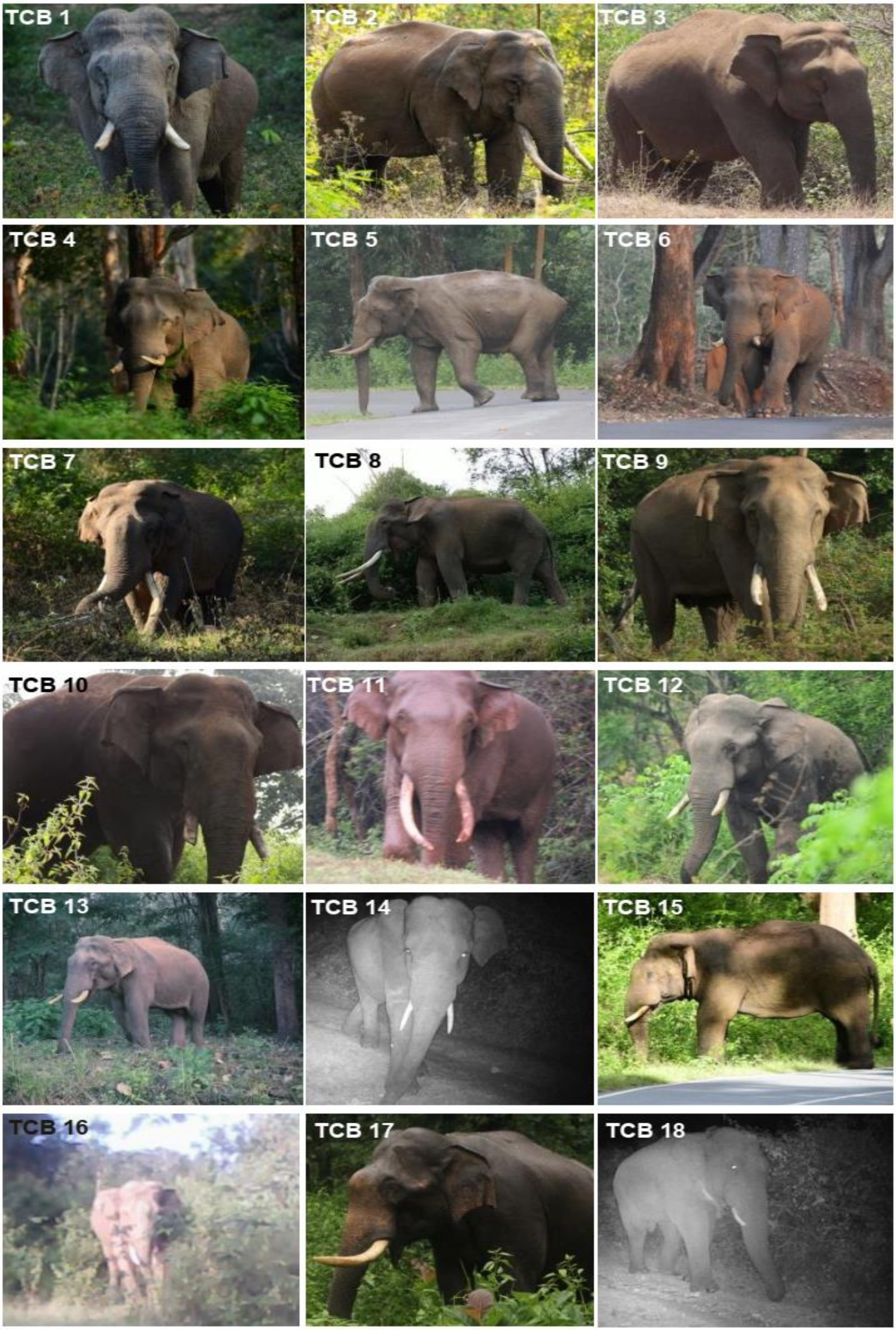

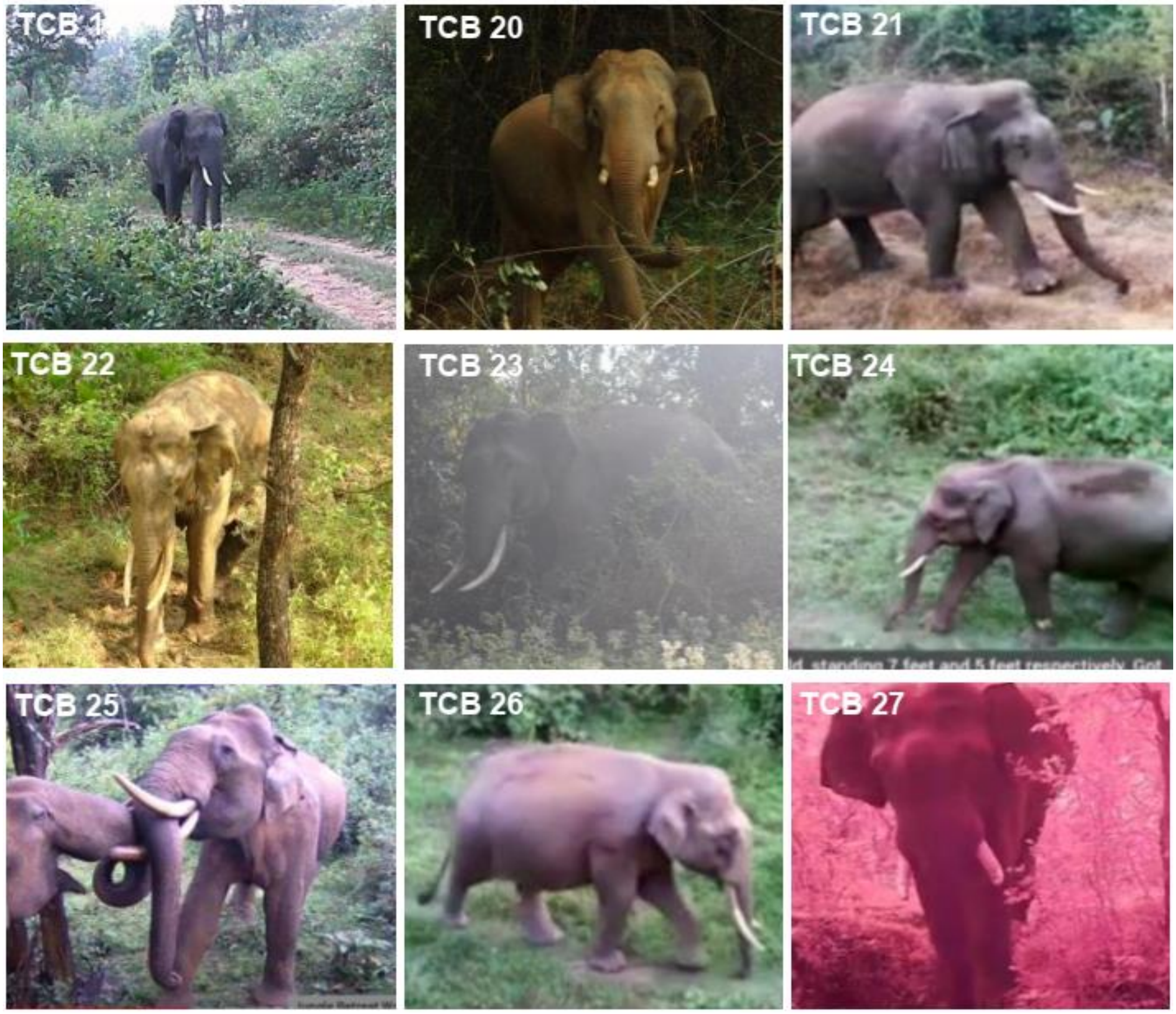
TCB 1
TCB 20
TCB 21
TCB 22
TCB 23
TCB 24
TCB 25
TCB 26
TCB 27

**Table S1.** Details of elephant sightings and behaviourin the Thirunelli-Kudrakote 'elephant corridor'

| Photo ID | Number of photographs | Approximate Age | Crop-raiding in the Corridor Area (Regular/Rare/No) | Musth period | Fidelity (High: >3 years, Low: ≤3 years) | Presence of External Body Scars/Injuries (Yes/No) | Notes |
|---|---|---|---|---|---|---|---|
| TCB1 | 60 | 30-35 | Regular | Sep-Nov | High | Yes | This individual is mostly found in the corridor, particularly in the area between Thirunelli and Kotayur village, but is rarely seen in the Tholpetty Range. During musth, he moves to the Brahmagiri hills. He is a persistent crop-raider in the area, displaying boldness but showing less aggression towards people. |
| TCB2 | 61 | 20-25 | Regular | June-August | High | Yes | This individual is observed throughout the corridor and Tholpetty Range. He is a persistent crop-raider, frequently targeting crops in Irumbupalam, Tholpetty Range, and along the corridor. Notably, he has not been recorded in Brahmagiri. He is a bold individual and exhibits aggressive behavior towards people. |
| TCB3 | 17 | 20-25 | Rare | Oct-Dec | High | Yes | Regularly seen along the corridor and in the north and southwestern parts of the Tholpetty Range during musth and non-musth season. He has not been recorded in Brahmagiri. This individual is bold and aggressive towards people. |
| TCB 4 | 17 | 25-30 | NA | Oct-Dec | High | Yes | This individual has not been observed raiding crops in the corridor area. Every year, he arrives in the corridor and Tholpetty area just before entering musth and remains there |

| | | | | | | | |
|---|---|---|---|---|---|---|---|
| | | | | | | | throughout the full musth period. After musth, he moves to other areas. Despite having external body injuries on his legs and flanks, he has not been involved in crop-raiding within the corridor. He is bold and less aggressive towards people. He has not been recorded in Brahmagiri. |
| TCB 5 | 42 | 30-35 | Regular | Oct-Nov | High | Yes | Commonly seen in the corridor and Tholpetty during both musth and non-musth seasons. He is bold and aggressive towards people. He is one of the most injured individual in the area due to negative interaction with people. He has not been recorded in Brahmagiri. |
| TCB 6 | 7 | 30-35 | NA | Oct-Dec | Low | NA | Observed him in the corridor and Tholpetty Range during musth. During non-musth, he is found in the Chedelath Range and Bavali, where he raid crops. He has not been recorded in Brahmagiri. |
| TCB 7 | 22 | 30-35 | No | Nov-Jan | High | NA | As one of the largest males in the area, he utilizes corridors during both musth and non-musth seasons. He tends to be elusive and is challenging to spot in open areas like TCB 1, TCB 2, or TCB 5, except during musth. Additionally, he has not been recorded in Brahmagiri. |
| TCB 8 | 16 | 25-30 | Rare | NA | High | NA | Recorded in the corridor and Tholpetty area, he is a typically elusive individual, seldom seen along the main road. He was observed raiding crops with TCB2 and has not been recorded in Brahmagiri. |
| TCB 9 | 5 | 35-40 | NA | NA | Low | NA | A large and elderly individual often seen in the company of TCB1 and frequently sighted in the Tholpetty Range. Notably, he |

| | | | | | | | |
|---|---|---|---|---|---|---|---|
| | | | | | | | has not been recorded in the Brahmagiri hills. His presence on the Wayanad side during the paddy and jackfruit season suggests involvement in crop raiding. |
| TCB 10 | 3 | 30-35 | NA | NA | Low | NA | During the paddy and jackfruit season, he is frequently found in the corridor, indicating a probable involvement in crop-raiding activities. |
| TCB 11 | 5 | 25-30 | NA | NA | Low | NA | |
| TCB 12 | 8 | 10-15 | Regular | NA | High | NA | Regularly engaged in crop-raiding within the northern parts of the Tholpetty Range and corridor, he is exclusively observed on the eastern side of the corridor. This individual is known for his aggressive behavior and is frequently spotted in the tourism zone of the Tholpetty Range. |
| TCB 13 | 19 | 25-30 | NA | NA | High | NA | He is known for crop-raiding behavior and is seldom spotted in open areas during daylight hours. |
| TCB 14 | 11 | 8-12 | NA | NA | NA | NA | |
| TCB 15 | 11 | 30-40 | NA | Oct-Dec | Low | NA | He was sighted in the corridor on only two occasions, both during musth periods. |
| TCB 16 | 4 | 15-25 | NA | NA | High | NA | |
| TCB 17 | 2 | 30-40 | NA | NA | High | NA | He was recorded in Tholpetty and the corridor area during the monsoon season but disappeared afterward. |
| TCB 18 | 7 | 15-20 | NA | NA | Low | Yes | He was tranquilized and relocated from Hassan district in Karnataka to Bandipur Tiger Reserve, approximately 130 km away. Initially photographed in the corridor on January 26, 2024 wearing a radio collar, he later |

| | | | | | | | |
|---|---|---|---|---|---|---|---|
| | | | | | | | moved to Mananthavady town in Wayanad district where he died due to improper management. |
| TCB 19 | 2 | 5-10 | NA | NA | Low | NA | |
| TCB 20 | 2 | 10-20 | NA | NA | Low | NA | |
| TCB 21 | 1 | 30-35 | NA | NA | Low | NA | |
| TCB 22 | 2 | 20-25 | NA | NA | Low | NA | |
| TCB 23 | 1 | 20-25 | NA | NA | Low | NA | |
| TCB 24 | 2 | 5-10 | NA | NA | Low | NA | |
| TCB 25 | 1 | 5-10 | NA | NA | Low | NA | |
| TCB 26 | 1 | 10-20 | NA | NA | Low | NA | |
| TCB 27 | 1 | 10-20 | NA | NA | Low | NA | |

26
